# Complex Models of Sequence Evolution Require Accurate Estimators as Exemplified with the Invariable Site plus Gamma Model

**DOI:** 10.1101/185652

**Authors:** Lam-Tung Nguyen, Arndt von Haeseler, Bui Quang Minh

## Abstract

The invariable site plus Γ model is widely used to model rate heterogeneity among alignment sites in maximum likelihood and Bayesian phylogenetic analyses. The proof that the invariable site plus continuous Γ model is identifiable (model parameters can be inferred correctly given enough data) has increased the creditability of its application to phylogeny reconstruction. However, most phylogenetic software implement the invariable site plus discrete Γ model, whose identifiability is likely but unproven. How well the parameters of the invariable site plus discrete Γ model are estimated is still disputed. Especially the correlation of the fraction of invariable sites with the fractions of sites with a slow evolutionary rate is discussed as being problematic. We show that optimization heuristics as implemented in frequently used phylogenetic software cannot always reliably estimate the shape parameter, the proportion of invariable sites and the tree length. Here, we propose an improved optimization heuristic that accurately estimates the three parameters. While research efforts mainly focus on tree search methods, our results signify the equal importance of verifying and developing effective estimation methods for complex models of sequence evolution.

---

In model based phylogenetic analysis, the invariable site plus Γ model (Yang 1994; Gu et al. 1995), hereafter referred to as I+Γ, is widely used to model rate heterogeneity among sites, because it often fits the data better than the Γ model or the invariable-sites model alone (Sullivan and Swofford 1997). Thus, the I+Γ model is frequently selected by MODELTEST (Posada and Crandall 1998). The I+Γ model has two parameters: the proportion of invariable sites *p*_inv_ (0 ≤ *p*_inv_ < 1) and the shape parameter *α* (>0) of the Γ distribution. A small *α* (<1) indicates strong rate heterogeneity, whereas a large *α* (>1) corresponds to weak rate heterogeneity. Under certain conditions *p*_inv_ and *α* compete with each other for the same phylogenetic signal. For example, *α* ≤ 1 already accounts for sites with low rates; that interferes with *p*_inv_ and causes a correlation between the parameters making reliable estimation of those parameters difficult (Sullivan et al. 1999; Mayrose et al. 2005). Despite this interference, it has been shown that the I+ continuous Γ model is identifiable for “*all but members of the F81 family of rate matrices on any phylogeny with more than two distinct interspecies distances*” (Rogers 2001; Allman and Rhodes 2008; Chai and Housworth 2011). Since the I+ continuous Γ model is identifiable, reliable parameter estimation for this model should be possible for sufficiently long multiple sequence alignments.

However, most phylogenetic software only implement the I+ discrete Γ (Yang 1994) model as an approximation to the continuous Γ model because of its computational efficiency. The discussed competition between *p*_inv_ and *α* is based on the analysis of the discrete Γ-distribution. The results have led to the suggestion to discourage the use of the I+ discrete Γ model (Yang 2006; Jia et al. 2014; Stamatakis 2014).

On the other hand, the identifiability of the I+ discrete Γ model is likely, but unproven (Chai and Housworth 2011) and it is unclear how accurately popular phylogenetic software estimate parameters of the I+ discrete Γ model.

Thus, we used simulations to assess the accuracy of the I+ discrete Γ estimators implemented in three maximum likelihood (ML) phylogenetic software: RAxML (Stamatakis 2014), PhyML (Guindon et al. 2010), IQ-TREE (Nguyen et al. 2015) and one Bayesian inference program MrBayes (Ronquist et al. 2012). More precisely, we simulated 100,000-bp long alignments along three balanced trees of 6, 24 and 96 taxa. The lengths of the alignments ensure the recovery of the correct tree topology. The three trees have uniform branch lengths of 0.1 substitutions per site except for one internal branch on the 6-taxon tree whose length equals 0.2 to allow for 3 distinct distances between the sequences as required for identifiability in the continuous case (Chai and Housworth 2011). We assumed the K2P model (Kimura 1980) with a transition/transversion ratio of 2.0 and the rate heterogeneity model I+ discrete Γ with four rate categories. For each tree and each pair (*p*_inv_, *α*) ∈ {0.0, 0.1, …, 0.9} × {0.1,0.5,1.0} we simulated 100 alignments using Seq-Gen (Rambaut and Grassly 1997). We used RAxML version 8.2.2, PhyML version 20141029, IQ-TREE version 1.3.7 and MrBayes version 3.2.6 compiled with the BEAGLE library (Ayres et al. 2012) to infer the invariable proportion, the shape parameter and the tree length from the simulated alignments. For RAxML, PhyML and IQ-TREE we used the default options.

For MrBayes we used the default priors, i.e., uniform distribution within interval [0,1] for *p*_inv_, exponential distribution with mean 1.0 for *α*, non-clocklike uniform Dirichlet distribution for branch lengths and *Γ* distribution with mean of 10 for tree lengths (Unconstrained:GammaDir(1.0,0.1,1.0,1.0)). The sequential version of MrBayes was run with four chains (one hot and three cold chains) and one million MCMC generations. 1,489 (16.5%) non-convergent MrBayes runs, where the effective sample sizes (ESS) on *p*_inv_, *α*, or tree lengths are smaller than 100, were repeated with five million generations. However, 52 of the extended re-runs were stopped after four weeks without completing all five million generations. We note that 207 of the 1,489 re-runs still did not converge. MrBayes estimates are then summarized as the mean of the posterior distribution with a default burn-in of 25%.

## Current Phylogenetic Programs Do Not Produce Accurate Estimates for the I+ Discrete Γ Model

Figure 1 displays the averages 
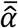
 of the estimated shape parameter *α*, the averages 
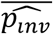
 of the estimated invariable fraction *p*_inv_ and the average 
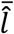
 of the estimated tree length *l* produced by PhyML, RAxML, IQ-TREE and MrBayes for the 100 alignments simulated from each parameter combinations. A program is called *accurate* if the estimated averages 
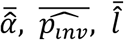
 deviate no more than 10% from the true values (depicted in green in Fig. 1).

**Figure 1.**
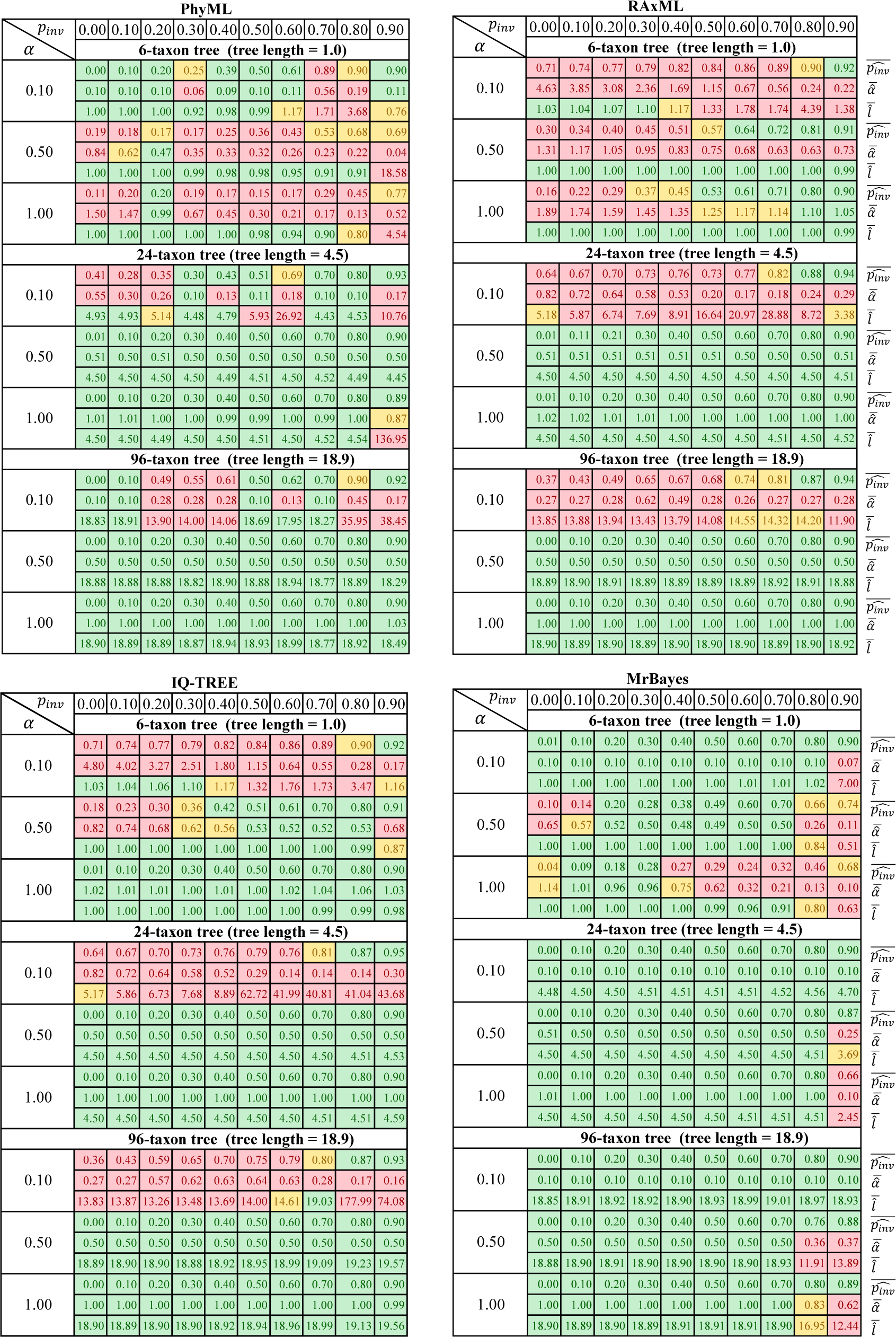
The averages 
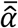
 of the estimated shape parameter *α*, the averages 
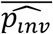
 of the estimated invariable fraction *p*_inv_ and the average 
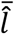
 of the estimated tree length *l* produced by PhyML, RAxML, IQ-TREE and MrBayes for the 100 alignments simulated from each parameter combinations. The averages are colored according to their differences from the true values: red (more than 25% deviation), yellow (10% to 25% deviation) and green (less than 10% deviation). For *p*_inv_ = 0.0 the estimated 
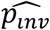
 is green if 
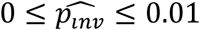
, yellow if 
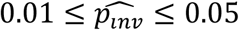
 and red if 
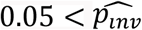
.

None of the tested programs estimated all parameter combinations accurately. The problem is especially pronounced for the 6-taxon alignments. For extreme rate heterogeneity (*α* = 0.1) MrBayes and PhyML recovered the true *α*, *p*_inv_ and *α* for 9/10 and 5/10 parameter combinations respectively, whereas the average estimates from IQ-TREE and RAxML were inaccurate. For strong rate heterogeneity (*α* = 0.5), the degrees of inaccuracy observed among all programs differ unsystematically. On the one hand, IQ-TREE and MrBayes accurately estimated the parameters in four and six settings. On the other hand, RAxML and PhyML could not estimate accurately the three parameters for any of the ten parameter-combinations. For medium rate variation (*α* = 1.0) only IQ-TREE produced the accurate estimates for all settings. All other programs exhibited varying degrees of inaccuracy.

For the 24‐ and 96-taxon alignments we observed an increase in the number of accurate estimates for all programs. These results corroborate a previous study (Sullivan et al. 1999) showing that increased taxon sampling leads to more reliable estimates. However, under extreme rate heterogeneity (*α* = 0.1), only MrBayes estimated all parameter sets accurately. We note that our measure of accuracy correlates well with the Bayesian coverage probabilities, the frequency with which true parameter values are included in the 95% credible interval of the estimates (Supplementary Fig. S1). 207 (2.3%) non-convergent MrBayes runs (effective sample size of *α* or *p*_inv_ are smaller than 100) partly overlap with cases where MrBayes was not accurate for 6-taxon simulations (*α* = 0.1 and *p*_inv_ = 0.9; *α* = 0.5 and *p*_inv_ ≤ 0.5; *α* = 1.0 and 0.*2* ≤ *p*_inv_ ≤ 0.5). Hence, non-convergence is a predictor of difficult settings but does not fully explain the inaccuracy of MrBayes (Fig. 1).

We also observed that inaccurate estimates of *α* and *p*_inv_ could sometimes lead to tree lengths that substantially deviate from the simulated lengths. For instance, for the 96-taxon alignments simulated with *α* = 0.1 and *p*_inv_ = 0.8 (expected tree length = 18.9) IQ-TREE estimated an average tree length of 177.0 that is nine times longer than the simulated tree length. The other programs also sometimes produced tree lengths that were considerably longer than the simulated ones.

In terms of computing times PhyML, RAxML and IQ-TREE needed for all analyses 62,441, 12,563 and 7,675 CPU hours, respectively. MrBayes needed 740,681 CPU hours to complete one million MCMC generations, thus it is 96.5 times slower than the fastest ML program. We note that this is only a lower-bound for the effective time one needs to wait for MrBayes results because 16.5% of MrBayes runs did not converge after one million MCMC generations. These runs were repeated with five times more generations, that led to significantly more computations.

## Multiple Local Optima on the Likelihood Surface Cause Inaccuracy

Because the tested programs performed quite differently with respect to the accuracy of parameter estimation, the number of taxa cannot be the only explanation. We suspected that the optimization heuristics as implemented in these programs drive the accuracy. Examining the likelihood surfaces for many simulated alignments revealed a common feature that the parameter space has two distinct peaks of high log-likelihoods (Fig. 2): one global close to the true parameters and one suboptimal peak with slightly lower log-likelihood (*ΔLNL* = −27 in this example) separated by a flat valley from the true parameters. In this particular instance MrBayes and PhyML found the true parameters whereas RAxML and IQ-TREE were trapped in the local maximum (not necessarily the case for other instances). In fact, whether the global or local optimum is detected depends on the starting values of the numerical optimization routines.

**Figure 2.**
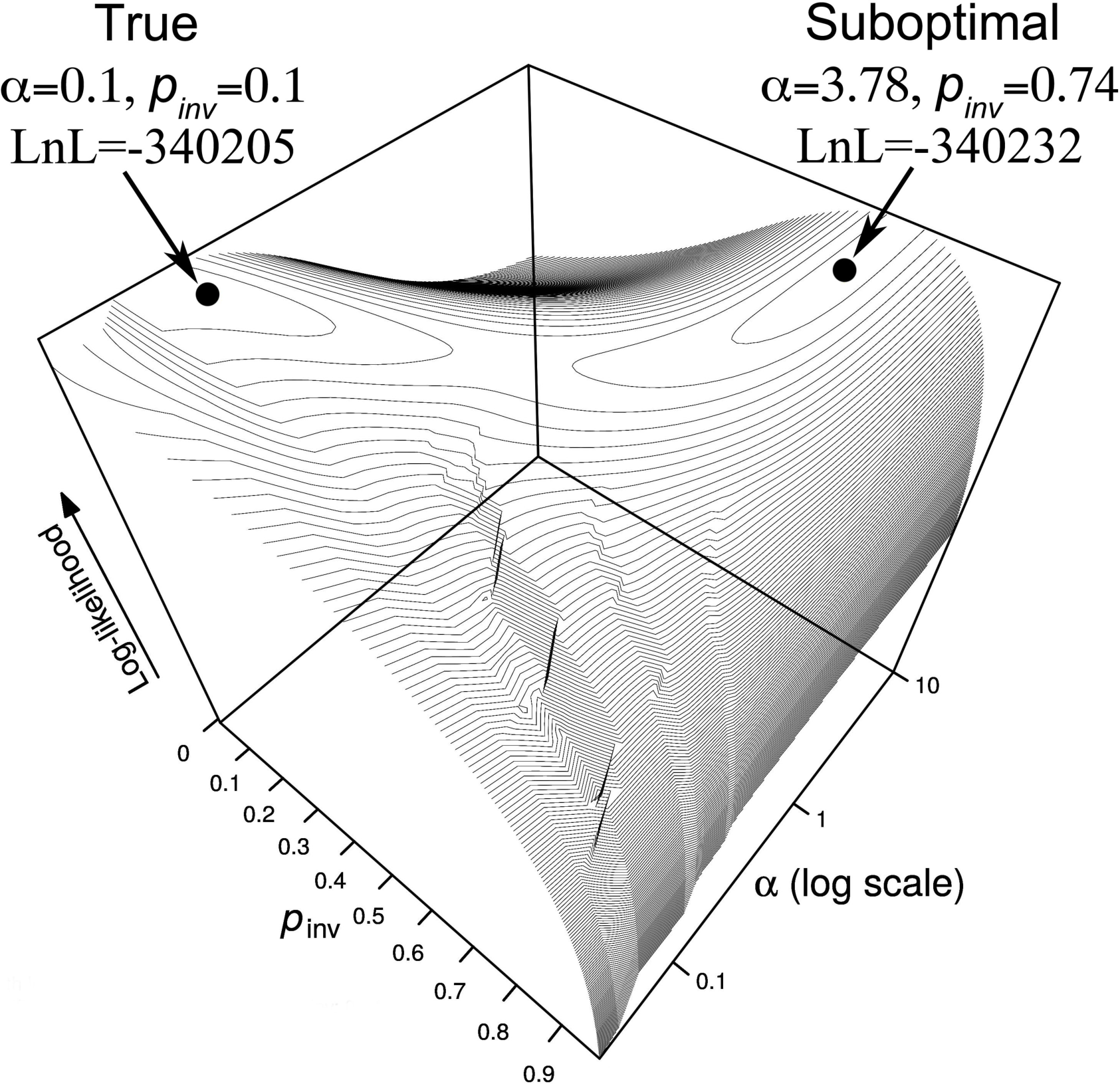
The likelihood surface for one simulated alignment as a function of *α* and *p*inv.

To summarize, we compared for each simulated alignment the log-likelihoods of the estimates with the log-likelihoods obtained for the true parameters. Table 1 show how often the true parameter combination produced a higher likelihood than the inferred parameters from MrBayes, PhyML, RAxML and IQ-TREE. These fractions are particularly high for the 6-taxon tree and for the ML inference programs. Most ML phylogenetic programs use general-purpose numerical methods to find *α* and *p*_inv_ (e.g., Brent 1973). These methods are obviously not well adapted to the complex likelihood surface (Fig. 2) and explain the poor overall performance of the ML programs (Fig. 1).

**Table 1.** Percentage of alignments where the true simulation parameters result in higher log-likelihoods than the inferred parameters from the programs for three simulation scenarios (6‐, 24‐ and 96-taxon trees).

| Program | 6-taxon tree | 24-taxon tree | 96-taxon tree |
| --- | --- | --- | --- |
| MrBayes | 34.7 % | 6.6 % | 5.6 % |
| PhyML | 60.2 % | 21.9 % | 45.0 % |
| RAxML | 89.2 % | 37.7 % | 49.5 % |
| IQ-TREE | 36.0 % | 34.1 % | 44.0 % |

## Effective Optimization Heuristic Produces Accurate Estimates

As remedy, we propose an alternative optimization heuristic which employs the EM algorithm (Dempster et al. 1977) to estimate *p*_inv_. We assume a discrete Γ distribution with *k* rate categories. Under the I+ discrete Γ model, the site rates follow a discrete mixture model consisting of *k* + 1 categories with rates *r*_0_, …, *r_k_*, where *r*_0_ = 0 represents invariable sites and *r_i_* > 0 (*i* = 1, ‥, *k*) are the *k* rates determined from the shape parameter *α* of the discrete Γ distribution (Yang 1994). Given a tree topology, the optimization heuristic does the following:

1. Choose initial values for α and *p*_inv_.
2. Optimize branch lengths by the Newton-Raphson method.
3. Optimize substitution model parameters by the BFGS method.
4. For each alignment site *D_i_* compute its posterior probability of being invariable (1 ≤ *i* ≤ *n*, where *n* is the number of alignment sites):

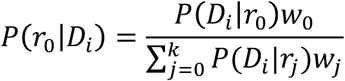

where *P*(*D_i_*|*r_j_* is the likelihood of site *D_i_* having rate *r_j_* and *w*_0_ = *p*_inv_, 
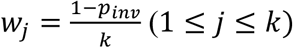
.
5. Update 
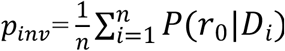
.
6. Optimize *α* by the Brent method.
7. If the log-likelihood improvement is greater than a predefined *ε* value, go back to step 2. Otherwise, stop the parameter optimization.

Step 4 and 5 correspond to the E‐ and M-step of the EM algorithm respectively. To avoid being stuck in local optima, we repeat this optimization procedure from ten starting values of *p*_inv_ evenly spaced between 0 and the fraction of constant sites observed in the alignment. The initial value of *α* is always set to 1.0.

We implemented the new optimization heuristic in IQ-TREE now called IQ-TREE-EM (IQ-TREE version 1.4.3) and repeated the previous simulations. Figure 3 shows that IQ-TREE-EM successfully recovered the true parameters for all but one parameter combination (6-taxon, *α* = 0.5 and *p*_inv_ = 0.0) where the average estimates 
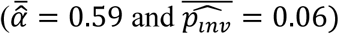
 slightly deviated from the true values.

**Figure 3.**
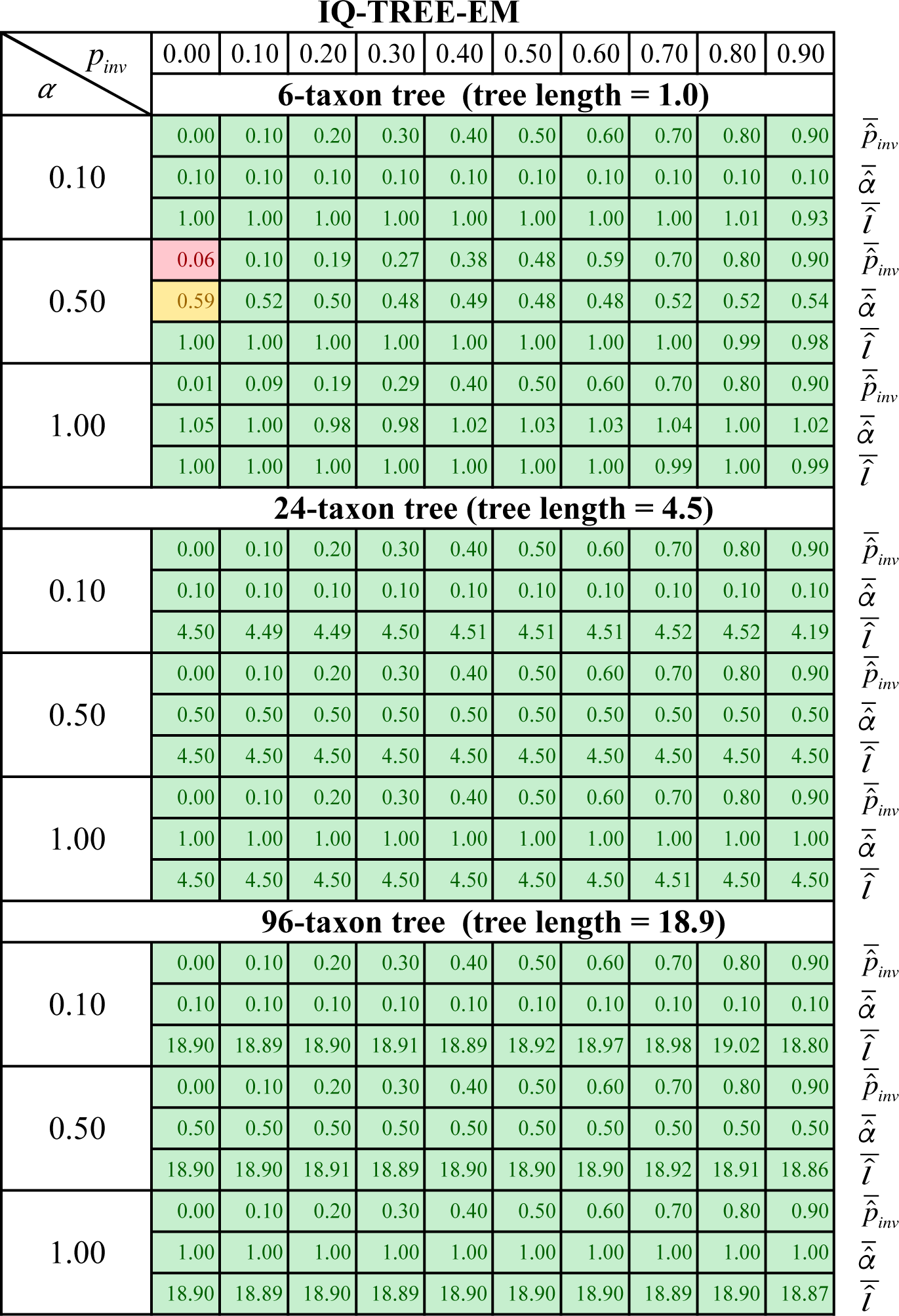
The averages 
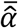
 of the estimated shape parameter *α*, the averages 
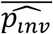
 of the estimated invariable fraction *p*_inv_ and the average 
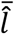
 of the estimated tree length *l* produced by IQ-TREE-EM for the 100 alignments simulated from each parameter combinations. The color code is explained in Figure 1.

Also, the percentage of instances where the estimated log-likelihoods were lower than the log-likelihood for the true parameters dropped considerably (0.06% 6-taxon tree, 0.0% 24-taxon tree, 0.03% 96-taxon tree; compare also with Table 1).

This increase in accuracy comes at the cost of an increased total computing time by a factor of 1.3 compared to IQ-TREE.

Thus, we conclude that the inaccurate parameter estimation of the I+ discrete Γ shown for the tested phylogenetic programs is caused by ineffective optimization methods.

## Impact On Real Data

To investigate the impact of accuracy on real data for ML estimates, we analyzed 70 DNA and 45 protein TreeBase alignments (Nguyen et al. 2015). We applied the GTR+I+Γ4 and LG+I+Γ4 models for DNA and protein data, respectively. Among 115 alignments, we detected 15 (5 DNA and 10 protein) alignments where the estimated *α* and *p*_inv_ by PhyML, RAxML or IQTREE deviated more than 10% from those by IQ-TREE-EM (Fig. 4; Supplementary Table S1). The estimates by PhyML and IQ-TREE deviated from those by IQ-TREE-EM only for one and two alignments, respectively. However, RAxML estimated *α* and *p*_inv_ dramatically different from IQ-TREE-EM, PhyML and IQ-TREE for all 15 alignments. Interestingly, RAxML systematically overestimated *α* and *p*_inv_ for all 5 DNA and underestimated them for all 10 protein alignments (*p*_inv_ sometimes very close to zero).

**Figure 4.**
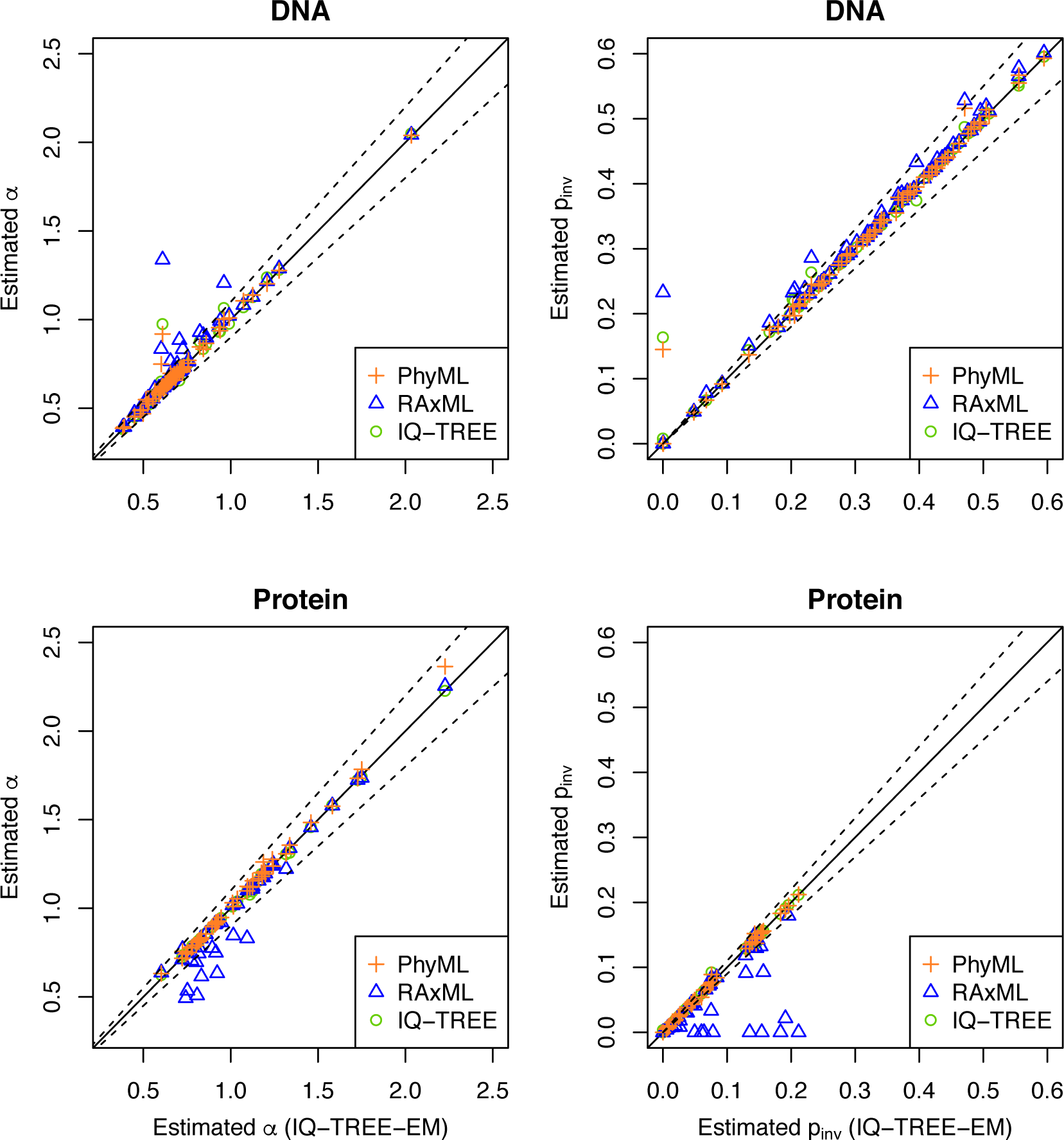
Estimation of *α* (left) and *p*_inv_ (right) for TreeBase alignments using IQ-TREE-EM (x-axis) and IQ-TREE (circle), PhyML (cross) and RAxML (triangle). Dashed lines show the boundaries of 10% deviation from the IQ-TREE-EM estimates. Points above the upper dashed lines indicate overestimation compared with IQ-TREE-EM, whereas points under the lower dashed lines indicate underestimation.

## Discussion

Our simulations revealed a major issue for parameter estimation of the I+ discrete Γ model as implemented in phylogenetic software. Despite using very long alignments, none of the tested programs recovered the true *α*, *p*_inv_ and tree length for all parameter combinations. Often, the estimates deviated heavily from the true values and different programs estimated different values for the same evolutionary parameters, although all programs inferred the true tree. Our further analysis of 115 TreeBase alignments showed that PhyML, IQ-TREE and IQ-TREE-EM estimates generally agree with each other except for two alignments. However, we identified 15 (13%) alignments where RAxML systematically overestimated *α* and *p*_inv_ for DNA and underestimated for protein, compared with other programs. The reasons for that behavior are unclear and deserve further analyses. While this result may not be extrapolated to other data sets, phylogenetic software should benefit from the more robust optimization described for IQ-TREE-EM.

We showed that the estimation heuristics implemented in popular phylogenetic programs causes such inaccurate estimates and the I+Γ model per se is not problematic. The relatively good performance of MrBayes is likely attributed to the Bayesian sampling of the parameter space but comes at the cost of excessive computing time.

With IQ-TREE-EM we provided an alternative optimization heuristic for ML methods that allows accurate estimation of the parameters for the I+ discrete Γ model. IQ-TREE-EM combines two optimization techniques: the multiple starting point strategy and the EM algorithm. We note that the EM algorithm alone will not achieve this accuracy (Supplementary Fig. S2). Therefore, while the former allows to escape local optima, the latter helps to speedup the optimization using analytical formula for *p*_inv_. This new approach effectively infers the true evolutionary parameters for long alignments. Thus, it is tempting to speculate that the GTR+I+ discrete Γ model is also identifiable as shown for the GTR+I+ continuous Γ model (Chai and Housworth 2011).

Our observations show that as models of sequence evolution become more and more complex (e.g., Dirichlet rate and other mixture models), tailored numerical optimization methods are necessary to achieve accurate estimates of evolutionary parameters. It is not enough to recover the true tree, if one wants to understand how evolutionary forces shaped contemporary genomes. The effect of wrong parameter estimates for the substitution model on the total tree length is sometimes dramatic (see Fig. 1). This may in turn bias downstream analysis such as divergence time dating, inference of site-specific evolutionary rates and ancestral sequence reconstruction, which are sensitive to the parameter estimates. Thus, one should critically scrutinize the heuristics implemented in popular programs. A more thorough evaluation of phylogenetic inference programs allowing for very complicated models of sequence evolution is necessary, but beyond the scope of this article.

Finally, we would like to point out that we only addressed the accurate computation of *p*_inv_ and *α* for the widely-used I+ discrete Γ model. We do not discuss the biological interpretation of *p*_inv_. The estimate of *p*_inv_ depends very much on the multiple sequence alignment at hand. *p*_inv_ may change if we enlarge the alignment. Thus, drawing an absolute conclusion from *p*_inv_ is in any case questionable.

## Supplementary Material

Data available from the Dryad Digital Repository: https://doi.org/10.5061/dryad.4j5c7.

## Acknowledgements

The authors would like to thank Heiko Schmidt for fruitful discussions, two anonymous reviewers, Fredrik Ronquist and Edward Susko for constructive and helpful comments on a earlier version of the manuscript. The computational results presented have been achieved using the Vienna Scientific Cluster 3 (VSC-3).

